# Description of a new member of the family *Erysipelotrichaceae: Clostridium fusiformis* sp. nov., isolated from healthy human feces

**DOI:** 10.1101/734715

**Authors:** Sudeep Ghimire, Supapit Wongkuna, Joy Scaria

**Author notes:** Address correspondence to: Joy Scaria.

## Abstract

A Gram-positive, non-motile, rod-shaped facultative anaerobic bacterial strain SG502^T^ was isolated from the healthy human fecal samples in Brookings, SD, USA. The comparison of the 16S rRNA gene placed the strain within the *Clostridium* cluster XVI, where, *Clostridium innocuum* ATCC 14501^T^, *Longicatena caecimuris* strain PG-426-CC-2, *Eubacterium dolichum* DSM 3991^T^ and *Eubacterium tortuosum* DSM 3987^T^ were its closest taxa with 95.15%, 94.49%, 93.28%, and 93.20% sequence identities respectively. The optimal growth temperature and pH for the strain SG502^T^ were 37°C and 7.0 respectively. Acetate was the major short-chain fatty acid product of the strain SG502^T^ when grown in BHI-M medium. The major cellular fatty acids produced by the strain SG502^T^ were C_18:1_ ω9c, C_18:0_ and C_16:0_. The DNA G+C content of the strain was 34.34 mol%. The average nucleotide identity of the genome of the strain SG502^T^ and its closest neighbor *C. innocuum* ATCC 14501^T^ was 63.48%. Based on the polyphasic analysis, the type strain SG502^T^ (=DSM 107282^T^), represents a novel species of the genus *Clostridium* for which the name *Clostridium fusiformis* sp. nov. is proposed.

## Introduction

The family *Erysipelotrichaceae* is an emerging group of bacteria originally described by Verbarg et al (1). The members of family *Erysipelotrichaceae* includes Gram-positive, filamentous rods and were originally described as facultative anaerobes but later amended by Tegtmeier *et al.* (2) to include obligate anaerobes. According to the updated LPSN list of valid bacteria (http://www.bacterio.net), current members of this family includes the following genera: *Allobaculum, Breznakia, Bulleidia, Catenibacterium, Canteisphaera, Coprobacillus, Dielma, Dubosiella, Eggerthia, Erysipelothrix, Faecalibaculum, Faecalicoccus, Faecalitalea, Holdemania, Kandleria, Longibacculum, Longicatena, Solobacterium*, and *Turicibacter.* Few species related to *Clostridium* and *Eubacterium* are also included within this family based on the 16S rRNA gene sequence similiarity and are considered as misclassified (3). Members of this family have low-G+C content and were previously recognized as the “walled relatives” of mycoplasma (4) and later classified under Clostridial cluster XVI (5). With major changes in the taxonomy of *Erysipelotrichaceae*, recently some members have been reclassified into new families *Coprobacillaceae* (5) and *Turicibacteraceae* (3) but the placement of cluster XVI is still debated.

The members of *Erysipelotrichaceae* have been isolated from the intestinal tracts of mammals (6-8) and insects (9) and are associated with host metabolism and inflammatory diseases (10, 11). Since there only few cultured species available from this family, their function in the gut ecosystem is not yet well understood (11).

Here, we describe the isolation, physiology and genome characteristics of a new member of the family *Erysipelotrichaceae* isolated from healthy human feces. The strain SG502^T^ clustered with clostridial cluster XVI and revealed genetic and phenotypic differences from *C. innocuum* strain the closest strain with valid taxonomy. Therefore, we propose the novel species *Clostridium fusiformis* sp. nov within family *Erysipelotrichaceae*.

### Isolation and Ecology

Strain SG502^T^ was isolated from the healthy human fecal sample. After transferring the fresh fecal samples into the anaerobic chamber (85% nitrogen, 10% hydrogen and 5% carbon dioxide) within 10 minutes of voiding, the sample was diluted 10 times with anaerobic PBS and stored with 18% DMSO in −80°C. The samples were cultured in modified BHI medium (BHI-M) containing 37g/L of BHI, 5 g/L of yeast extract, 1 ml of 1 mg/mL menadione, 0.3 g of L-cysteine, 1 mL of 0.25 mg/L of resazurin, 1 mL of 0.5 mg/mL hemin, 10 mL of vitamin and mineral mixture,1.7 mL of 30 mM acetic acid, 2 mL of 8 mM propionic acid, 2 mL of 4 mM butyric acid, 100 μl of 1 mM isovaleric acid, and 1% of pectin and inulin. After isolation, the strain was subjected to MALDI-ToF (Bruker, Germany). Since MALDI-TOF did not identify a species,, 16S rRNA gene sequencing was performed for species validation.

### Physiology and Chemotaxonomy

The strain was cultivated in BHI-M medium in anaerobic conditions at 37°C at pH 6.8±0.2 for morphological characterization. Bacterial colony characteristics were determined after streaking the bacteria on BHI-M agar plates followed by 48 hours of anaerobic incubation. Gram staining was performed using Gram staining kit (Difco) according to the manufacturer’s instruction. During the exponential growth of the bacterium, cell morphology was examined by scanning electron microscopy (SEM). SG502^T^ was grown separately under aerobic and anaerobic conditions to determine the aerotolerance. Further, the strain was grown at 4, 20, 30, 40 and 55°C to determine the range of growth under anaerobic conditions. The BHI-M media was adjusted to pH range of 4-9 with 0.1N HCl and 0.1N NaOH to determine the growth of the strain at different pH levels. BHI-M medium was supplemented with triphenyltetrazolium chloride (TTC) (12) to determine the motility of the strain in which red color indicated positive motility.

Individual cells of the strain SG502^T^ was gram-positive rod-shaped with 1.5×0.35μ in dimensions (Fig 1 and Table 1). The colonies appeared to be white, smooth and convex with entire edges. The strain SG502^T^ grew from a pH range of 6.0-7.5 with optimal growth at pH 7.0 at 37°C anaerobically. The bacterium could grow anaerobically over the temperature range of 25-45°C with optimal growth at 37°C.

**Table 1:**
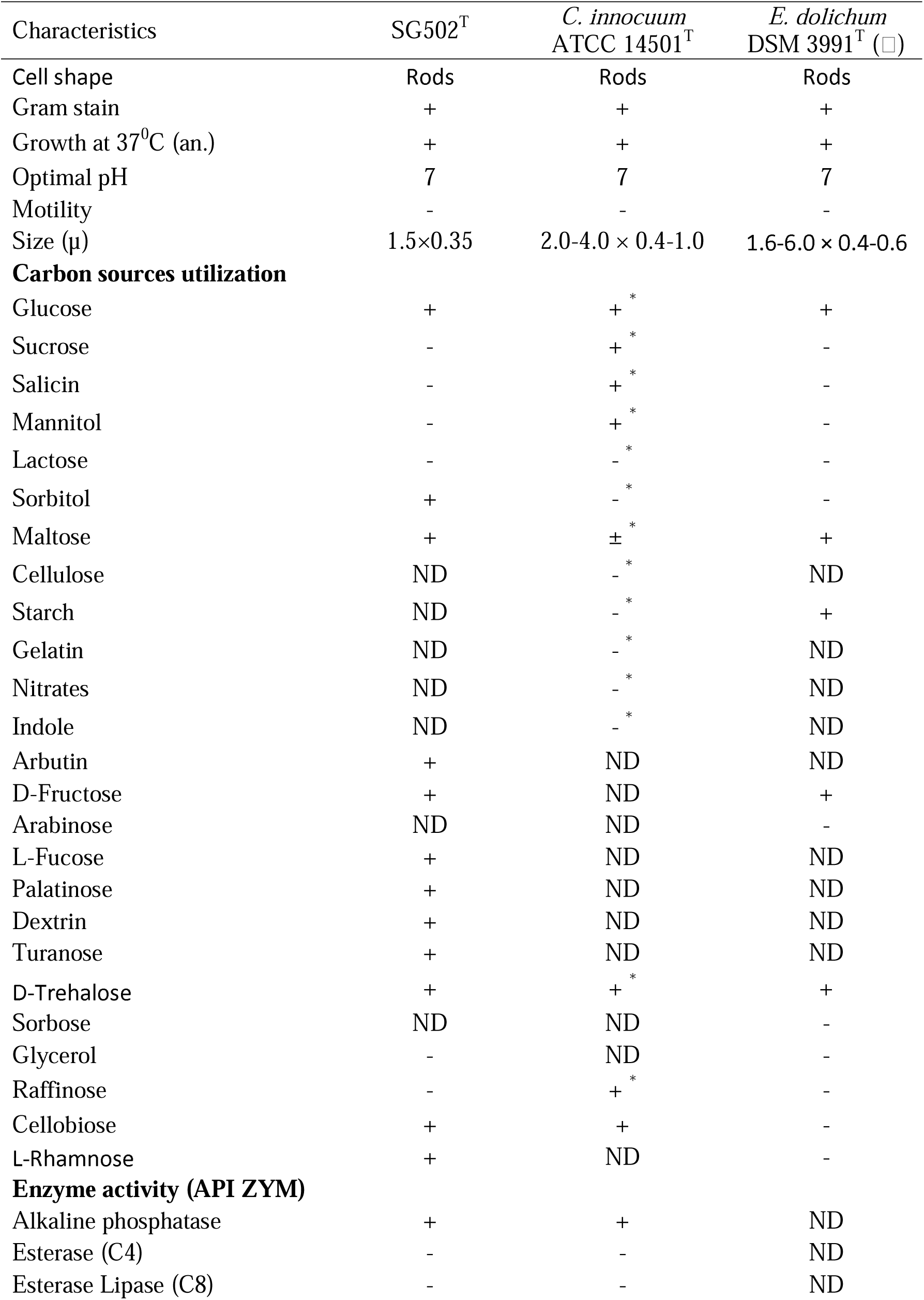
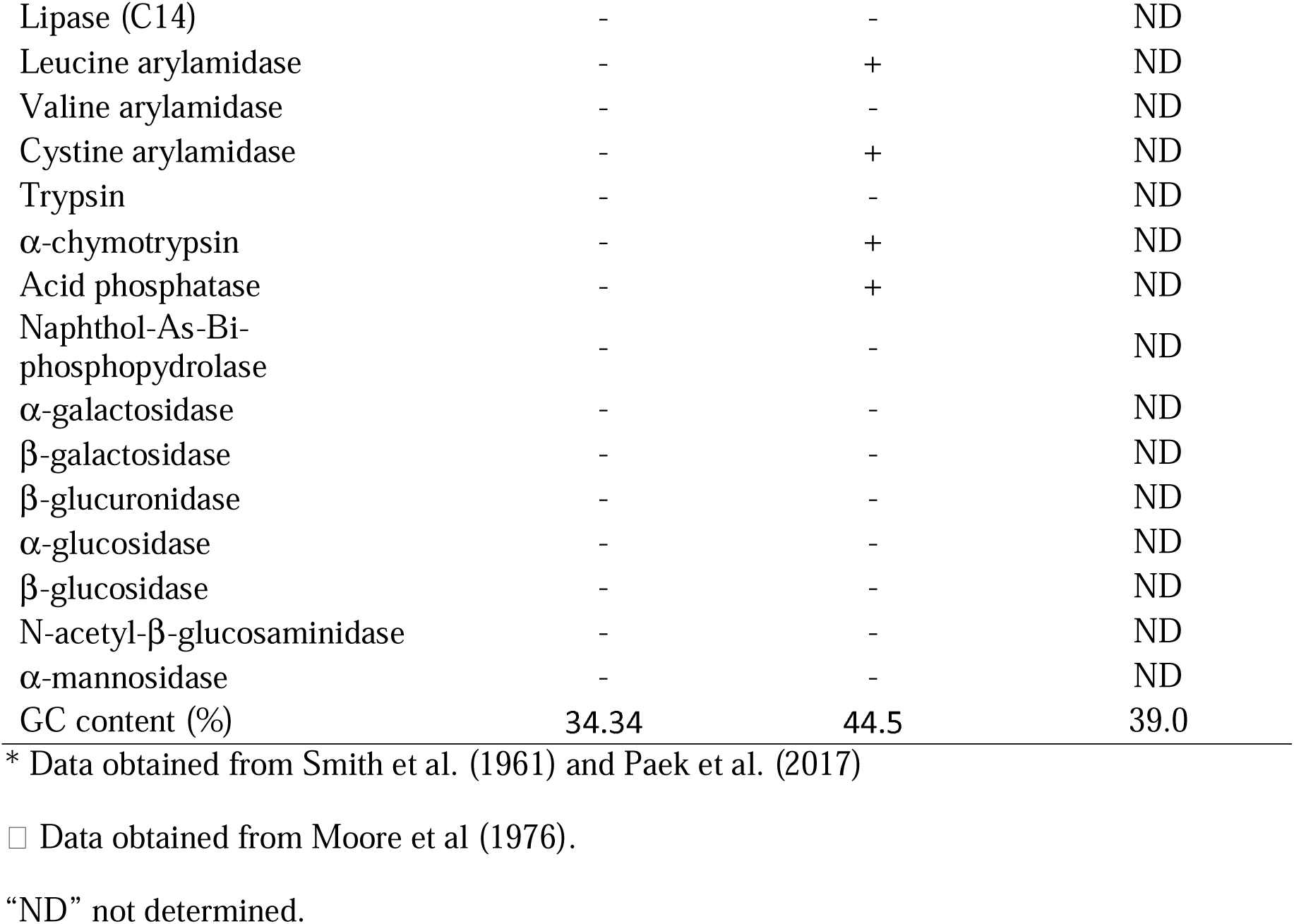
Differential biochemical features of the strain SG502^T^ and its closest phylogenetic neighbor *C. innocuum* ATCC 14501^T^ identified using API ZYM (bioMerieux, France).

**Fig. 1:**
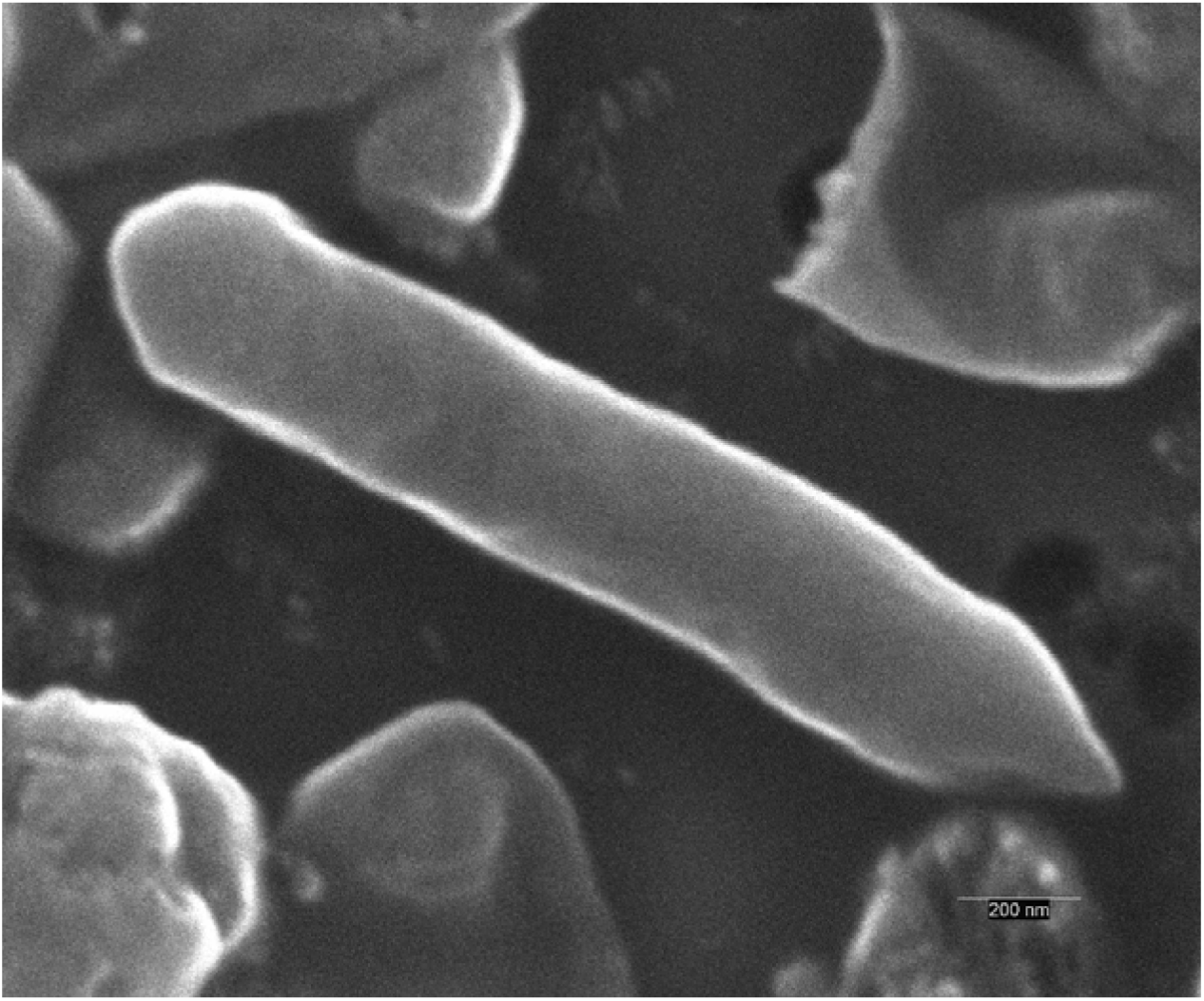
Scanning electron micrograph of strain SG502^T^. Cells were imaged after culturing in anaerobic conditions for 24 hours at 37°C in BHI-M medium. Bar, 200 nm.

The strain grew well in BHI-M broth under anaerobic conditions but under aerobic conditions, the growth was comparatively lower and prolonged confirming that the strain was a facultative anaerobe. The utilization of various carbon sources and enzymatic activities were determined using AN MicroPlate (Biolog) and API ZYM (bioMerieux) according to the manufacturer’s instructions. After growing the strain SG502^T^ and ATCC 14501^T^ in BHI-M medium at 37°C for 24 hours, cells were harvested for cellular fatty acid analysis. Fatty acids were extracted, purified, methylated and identified and analyzed using GC (Agilent 7890A) according to manufacturer’s instructions (MIDI) (13). Short-chain fatty acids were determined using gas chromatography after cells were grown in BHI-M medium. For SCFAs, the culture was maintained in 25% meta-phosphoric acid before collecting the supernatant for analysis to detect volatile fatty acids.

Based on the results obtained from Biolog AN plate, the strain SG502^T^ utilizes glucose, sorbitol, maltose, arbutin, D-fructose, L-fucose, palatinose, dextrin, turanose, D-trehalose, L-rhamnose, uridine, pyruvic acid methyl ester, pyruvic acid, 3-methyl-D-glucose, gentiobiose, maltotriose, ducitol, L-phenylalanine, α-ketovaleric acid, N-acetyl-D-glucosamine, N-acetyl-β-D-mannosamine, cellobiose, a-ketobutyric acid, D-galacturonic acid and N-acetyl-D-glucosamine. Also, the strain SG502^T^ assimilated sorbitol and maltose which were not utilized by ATCC 14501^T^. Furthemore, SG502^T^ was unable to utilize sucrose, salicin, mannitol, lactose, and raffinose when compared to ATCC 14501^T^ strain. Positive enzymatic activities for leucine arylamidase, cystine arylamidase, α-chymotripsin and acid phosphates were observed for ATCC 14501^T^ differentiating it from strain SG502^T^ (Table1). The major SCFAs identified for SG502^T^ was acetate. Low but detectable amounts of propionate and butyrate were produced by the strain SG502^T^.

### 16S rRNA phylogeny

DNA was isolated using E.Z.N.A bacterial DNA isolation kit (Omega Biotek) following the manufacturer’s instructions. The 16S rRNA gene was amplified using universal primer set 27F (5’-AGAGTTTGATCMTGGCTCAG-3’) and 1492R (5’-ACCTTGTTACGACTT-3’) and sequenced using a Sanger sequencing chemistry (ABI 3730XL; Applied Biosystems). The sequences were assembled using Genious 10.2.3. The 1338 bp of the 16S rRNA gene was obtained and searched against the NCBI 16S rRNA gene database for identification. The closest species identified were *Clostridium innocuum* strain B-3, *Longicatena caecimuris* strain PG-426-CC-2, *Eubacterium dolichum* and *Eubacterium tortuosum* with 95.15%, 94.49%, 93.28%, and 93.20% sequence identities respectively. The 16S rRNA gene sequence alignment and phylogenetic analysis were conducted using MEGA7 software (15). Initially, the sequences were aligned using CLUSTAL-W (16) and neighbor-joining (17) method was used to reconstruct the phylogenetic tree employing Kimura’s two-parameter model (18) with 1000 bootstraps. Based on the NJ phylogeny of the strain SG502^T^, the closest relative was found to be *Clostridium innocuum* ATCC 14501^T^, *Longicatena caecimuris* PG-426-CC-2, *Eubacterium dolichum* and *Eubacterium tortuosum* strains (Fig. 2). Currently, the cut off for the genus-level classification of the bacteria based on 16S rRNA gene is 94.5% identity (14). Since SG502^T^ was found to be 95.15% identical to *C. innocuum*, it was classified within genus *Clostridium*. With the differences in physiological, biochemical, and 16s rRNA sequence dissimiliarities of SG502^T^ with *C. innocuum*, we propose classifying it as a new species under genus *Clostridium*.

**Fig. 2:**
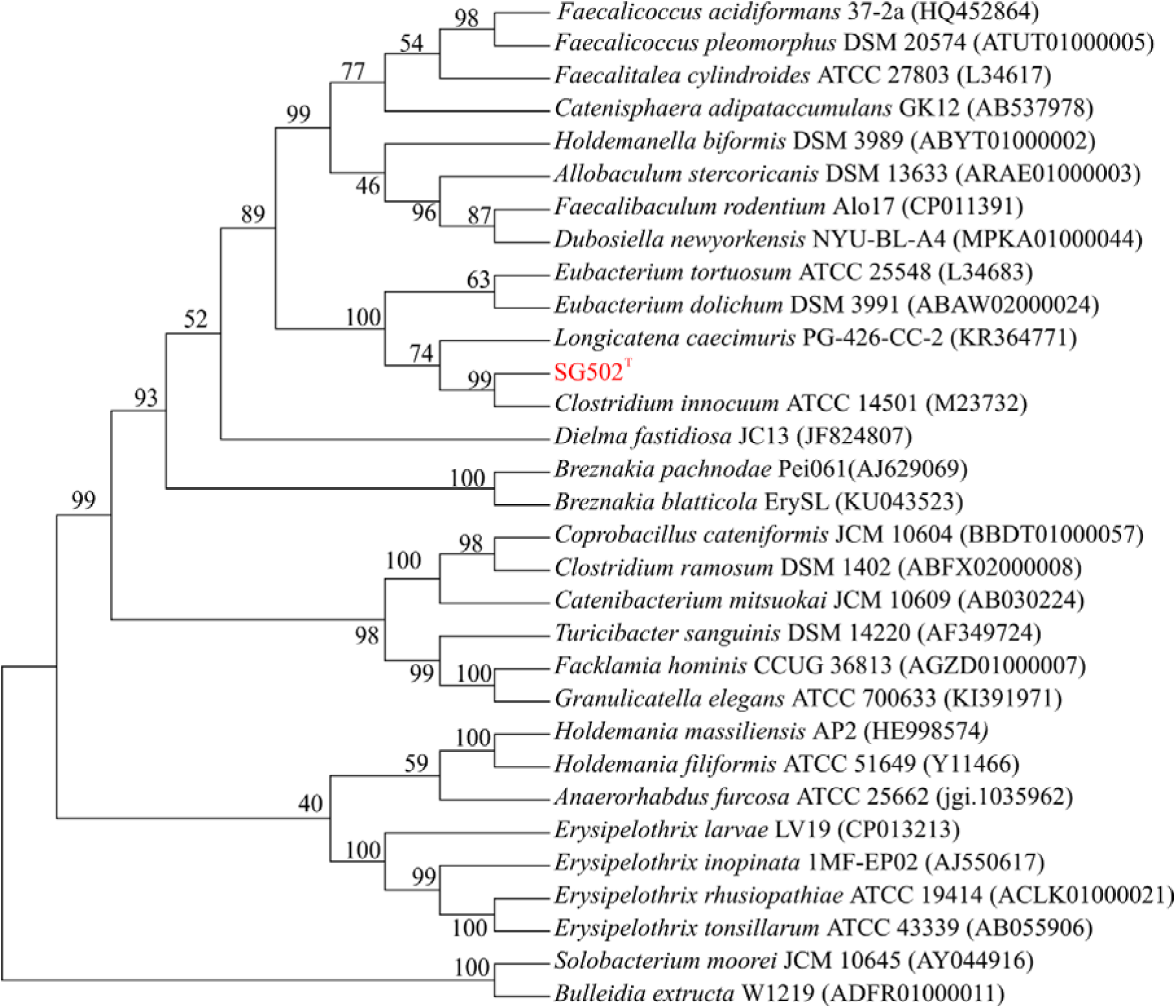
Neighbor-joining phylogenetic tree of 16S rRNA gene sequences of SG502^T^ with related species with a valid taxonomy. GenBank accession numbers of the 16S rRNA gene sequences are given in parentheses. Bootstrap values are shown as percentages at each node. The evolutionary history was inferred using the Neighbor-Joining method with 1000 bootstraps in MEGA7.

### Genome Features

The whole-genome sequencing of strain SW165 was performed using Illumina MiSeq using 2x 250 paired-end V2 chemistry. The reads were assembled using Unicycler that builds an initial assembly graph from short reads using the de novo assembler SPAdes 3.11.1 (19). The quality assessment for the assemblies was performed using QUAST (20). Genome annotation was performed using Prokka (21). The draft genome of the strain SG502 consisted of 146 contigs and was 2,387,606 bps long with 34.34 mol% G+C content. The total number of predicted coding sequences, tRNAs, rRNAs, and tmRNAs was 2348, 63, 2 and 1 respectively. The average nucleotide identity between the two type strains SG502^T^ and *C. innocuum* (NZ_AGYV00000000.1) was found to be 63.48% (22). This delineates the two strains from one another as the proposed cut off for ANI for a new species is 95-96% (23).

### Description of Clostridium sp. SG502^T^ sp. nov

#### Clostridium fusiformis

(Latin: “*fusus”*: spindle-like, referring to the shape of the bacterium)

The cells of the bacterium are anaerobic, gram-positive non-motile rods. The average size of the cell is 1.5×0.35 μ. Bacterial colonies on BHI-M agar are white, convex and entire approximately 0.1 cm in diameter. The optimum temperature and pH for the anaerobic growth are 37°C and 7.0 respectively. The strain SG502^T^ utilizes glucose, sorbitol, maltose, arbutin, D-fructose, L-fucose, palatinose, dextrin, turanose, D-trehalose, L-rhamnose, uridine, pyruvic acid methyl ester, pyruvic acid, 3-methyl-D-glucose, gentiobiose, maltotriose, ducitol, L-phenylalanine, a-ketovaleric acid, N-acetyl-D-glucosamine, N-acetyl-b-D-mannosamine, cellobiose, a-ketobutyric acid, D-galacturonic acid and N-acetyl-D-glucosamine. Positive enzymatic reactions were observed for alkaline phosphatase only. The primary short-chain fatty acid produced by the strain is acetate while small amounts of propionate and butyrate were also noted. The major cellular fatty acids of the strain SG502^T^ are C_18:1_ ω9c, C_18:0_ and C_16:0_. (summarized in Table 2). The type strain, SG502^T^ (=DSM 107282^T^), was isolated from a healthy human fecal sample. The DNA G+C content of the strain SG502^T^ is 34.34 mol%.

**Table 2:** Cellular fatty acid contents percentages of strain SG502^T^ compared to its phylogenetic neighbors *C. innocuum* ATCC 14501^T^ and *E. dolichum* DSM 3987^T^.

| Characteristics | SG502<sup>T</sup> | *C. innocuum*<br>ATCC 14501<sup>T</sup> | *E. dolichum* DSM<br>3987<sup>T</sup> (‡) |
| --- | --- | --- | --- |
| **Straight chain** |  |  |  |
| C<sub>10:0</sub> | 0.31 | 0.32 | 1.0 |
| C<sub>12:0</sub> | 1.74 | 3.28 | - |
| C<sub>14:0</sub> | 3.54 | 8.85 | 1.6 |
| C<sub>16:0</sub> | **14.70** | **23.70** | **22.8** |
| C<sub>16:0</sub> aldehyde | 0.42 | 2.78 | - |
| C<sub>17:0</sub> | 1.41 | - | 1.8 |
| C<sub>18:0</sub> | **22.55** | **10.56** | **17.1** |
| C<sub>18:0</sub> aldehyde | - | 1.16 | - |
| **Dimethyl acetal (DMA)** |  |  |  |
| C<sub>16:0</sub> DMA | 1.46 | 9.95 | - |
| C<sub>18:0</sub> DMA | 3.62 | 2.83 | - |
| **Unsaturated** |  |  |  |
| C<sub>16:1</sub> ω9c | 0.99 | 6.51 | - |
| C<sub>16:1</sub> ω7c | **6.47** | **10.59** | - |
| C<sub>16:1</sub> cis 9 DMA | - | - | - |
| C<sub>18:1</sub> ω9c | **29.82** | **14.64** | **33.9** |
| C<sub>18:1</sub> ω7c | **10.77** | **3.08** | - |
| C<sub>18:2</sub> cis 9,12 | - | - | 9.50 |
| C<sub>18:1</sub> cis 9 DMA | - | - | - |
| Summed Feature 5 | - | - | 6.5 |
| Summed Feature 8 | 0.54 | - | - |
| Summed Feature 10 | 10.77 | 3.08 | - |

(‡) Data from Paek et al. (2017)
Those fatty acids which were not separated using MIDI system were considered as summed features. Summed feature 5 contains C<sub>15:0</sub> DMA or C<sub>14:0</sub> 3-OH; summed feature 8 contains C<sub>17:1</sub> cis 9 or C<sub>17:2</sub> and summed feature contains C<sub>18:1</sub> c11/t9/t6 or UN17.83Q.

### Protologue

The GenBank BioProject ID number for the draft genome sequence of the strain SG502^T^ is PRJNA494608.

## Acknowledgments

The authors would like to thank Electron Microscopy Core Facility at the Bowling Green State University, Ohio, USA for assistance with scanning electron microscopy.

## ABBREVIATIONS

MALDI-ToF: Matrix Assisted Laser Desorption/Ionization-Time of Flight
ANI: Average Nucleotide Identity

